# Molecular diagnosis of orthopaedic device infection direct from sonication fluid by metagenomic sequencing

**DOI:** 10.1101/118026

**Authors:** Teresa L. Street, Nicholas D. Sanderson, Bridget L. Atkins, Andrew J. Brent, Kevin Cole, Dona Foster, Martin A. McNally, Sarah Oakley, Leon Peto, Adrian Taylor, Tim E.A. Peto, Derrick W. Crook, David W. Eyre

## Abstract

Culture of multiple periprosthetic tissue samples is the current gold-standard for microbiological diagnosis of prosthetic joint infections (PJI). Additional diagnostic information may be obtained through sonication fluid culture of explants. However, current techniques can have relatively low sensitivity, with prior antimicrobial therapy and infection by fastidious organisms influencing results. We assessed if metagenomic sequencing of complete bacterial DNA extracts obtained direct from sonication fluid can provide an alternative rapid and sensitive tool for diagnosis of PJI.

We compared metagenomic sequencing with standard aerobic and anaerobic culture in 97 sonication fluid samples from prosthetic joint and other orthopaedic device infections. Reads from Illumina MiSeq sequencing were taxonomically classified using Kraken. Using 50 samples (derivation set), we determined optimal thresholds for the number and proportion of bacterial reads required to identify an infection and validated our findings in 47 independent samples.

Compared to sonication fluid culture, the species-level sensitivity of metagenomic sequencing was 61/69(88%,95%CI 77-94%) (derivation samples 35/38[92%,79-98%]; validation 26/31[84%,66-95%]), and genus-level sensitivity was 64/69(93%,84-98%). Species-level specificity, adjusting for plausible fastidious causes of infection, species found in concurrently obtained tissue samples, and prior antibiotics, was 85/97(88%,79-93%) (derivation 43/50[86%,73-94%], validation 42/47[89%,77-96%]). High levels of human DNA contamination were seen despite use of laboratory methods to remove it. Rigorous laboratory good practice was required to prevent bacterial DNA contamination.

We demonstrate metagenomic sequencing can provide accurate diagnostic information in PJI. Our findings combined with increasing availability of portable, random-access sequencing technology offers the potential to translate metagenomic sequencing into a rapid diagnostic tool in PJI.

## Introduction

Prosthetic joint infections (PJI) are a devastating and difficult to treat complication of joint replacement surgery. Although the relative incidence of PJI is low (0.8% knee and 1.2% hip replacements across Europe) (1), given the increasing numbers of arthroplasties performed worldwide it is a significant healthcare burden and cause of expense. For individual patients, PJI often requires multiple surgeries, intensive, long-term antimicrobial therapy, and a prolonged period of rehabilitation. Fast, accurate and reliable diagnosis of PJI is necessary to inform treatment choices, particularly for antibiotic resistant organisms. Culture of multiple periprosthetic tissue (PPT) samples remains the gold-standard method of microbial detection (2–4). However, culture can be relatively insensitive with only 65% of causative bacteria detected in infections even when multiple PPT samples are collected (2, 5). Infections with fastidious organisms or where a patient has received prior antimicrobial treatment are often culture-negative.

Culture of sonication fluid from explanted prostheses may improve microbiological yield in PJI, by disrupting bacterial biofilm. Since sonication was first applied to explanted hip prostheses in 1998 (6) several clinical studies have reported improved sensitivity of sonication fluid culture over PPT culture for the diagnosis of hip, knee and shoulder PJI (7, 8), and sonication has been adopted by many centers, either alone or in combination with PPT culture. Additionally, several molecular assays have been investigated to improve the sensitivity of PJI diagnosis. PCR assays using DNA extracted from sonication fluid (9–12) have reported sensitivity ranging from 70% to 96%. However, this approach can only identify pathogens in a pre-defined multiplex panel, thus may miss atypical or rare pathogens not targeted in the assay design. Other studies identify pathogens by amplification and sequencing of the universal bacterial 16S ribosomal RNA gene (13, 14). A drawback of these methods is the potential for generating false-positive results from contaminating bacterial DNA.

The potential of high-throughput sequencing as a diagnostic tool for infectious diseases is widely recognized (15–17). Metagenomic sequencing offers the possibility to detect all DNA in a clinical sample, which can be compared to reference genome databases to identify pathogens. Additionally, a profile of common laboratory and kit contaminants can be generated from negative controls sequenced concurrently and accounted for (18, 19). In addition to diagnostic data, whole-genome sequencing can also simultaneously provide characterization of infection outbreaks (20, 21), tracking of transmission (22–24) and prediction of antimicrobial resistance (25–28). An advantage offered by sequencing is the speed at which it can deliver genetic information (29) compared to traditional microbiological culture and antimicrobial susceptibility testing, which can take days to weeks depending on the pathogen. By removing a culture step and sequencing directly from clinical samples the time taken to diagnosis can be reduced further (30) and pathogens not identified by conventional methods can be detected (31–33). Here, we investigated if metagenomic sequencing of complete bacterial DNA extracts obtained direct from sonication fluid can provide an alternative rapid and sensitive tool for diagnosis of PJI, without the need for a culture step.

## Materials and Methods

### Sample collection and processing

Intra-operative samples from the Nuffield Orthopaedic Centre (NOC) in Oxford University Hospitals (OUH), UK, between June 2013 and January 2017 were investigated. The NOC is a tertiary level specialist musculoskeletal hospital, including a dedicated Bone Infection Unit, undertaking approximately 200 revision arthroplasties annually. A subset of samples submitted were chosen at random following culture to provide a ratio of approximately 2:1 bacterial culture-positive samples to culture-negative samples.

Prosthetic joint implants and metalwork, received into the OUH microbiology laboratory following revision arthroplasty and operative management of other orthopaedic device related infection, were placed directly into single-use sterile polypropylene containers (Lock & Lock brand) and covered with between 10ml and 400ml of sterile 0.9% saline solution (Oxoid Ltd, Basingstoke, UK) depending on the size of the prosthesis/device, with sufficient fluid to cover at least 90% the prosthesis/device, up to a maximum of 400ml. Sonication was performed as described previously (7) with minor modifications. Briefly, the implant was vortexed for 30 seconds, subjected to sonication for 1 minute followed by additional vortexing for 30 seconds. Sonication was performed in a Bransonic 5510 ultrasonic water bath (Branson, Danbury, CT, USA) at a frequency of 40kHz. The resulting sonication fluid was plated in 0.1ml aliquots onto Columbia blood (CBA) and chocolate agar plates for aerobic incubation and CBA plates for anaerobic incubation. Aerobic incubation was performed at 35-37°C with 5% CO_2_ for up to 5 days. Anaerobic incubation was performed at 35-37°C for 10 days. All cultured microorganisms were identified by MALDI-TOF on a Microflex LT using Biotyper v.3.1 (Bruker Daltonics, Billerica, MA, USA). Samples were recorded as culture-positive where ≥50 CFU/ml were observed, and additionally at <50 CFU/ml for organisms considered highly pathogenic (including *Staphylococcus aureus* and Enterobacteriaceae.).

Periprosthetic tissue samples were also collected during surgery and processed by the microbiology laboratory. Briefly, BACTEC bottles were inoculated with 0.5ml of an inoculum generated by vortexing each tissue sample in 3ml 0.9% saline with Ballotini balls for 15 seconds. Bottles were incubated under aerobic (Plus Aerobic/F culture vials) and anaerobic (Lytic/10 Anaerobic/F culture vials) conditions in a BD BACTEC FX system (BD Biosciences, Sparks, MD, USA) for up to 10 days. Any bottles that flagged positive were subcultured onto agar plates and processed as described above to determine species.

### Bacterial DNA extraction from sonication fluid

Prior to DNA extraction sonication fluids were concentrated by centrifugation. 40ml of fluid was transferred to a sterile, disposable 50ml polypropylene tube and centrifuged at 15,000 × g in a Sorvall RC5C Plus centrifuge (SLA-1500 rotor with custom made inserts) for 1 hour at 16°C. Samples with <40ml starting volume of sonication fluid were made up to 40ml with the same saline used for sonication. All but approximately 1ml of the supernatant was discarded, and the pellet resuspended in this volume of fluid before being passed through a 5 μm syringe filter to deplete the number of human cells present, and therefore the amount of human DNA in the final extract. Bacterial cells passing through the filter were pelleted, washed with 1ml 0.9% saline and resuspended in 500μl molecular biology grade water before being mechanically lysed in Pathogen Lysis tubes (S) (Qiagen, Hilden, Germany) with a Fastprep-24 tissue homogeniser (MP Biomedicals, Santa Ana, CA, USA) (3 × 40 seconds at 6.5 m/s). DNA was extracted by ethanol precipitation, using GlycoBlue (Life Technologies, Paisley, UK) as a co-precipitant, and resuspended in 50μl 1 x Tris-EDTA (TE) buffer. DNA was purified using AMPure XP solid phase reversible immobilisation (SPRI) beads (Beckman Coulter, High Wycombe, UK) and eluted in 26μl TE buffer. DNA concentration was measured using a Qubit 2.0 fluorometer (Life Technologies, Paisley, UK). A subset of samples were treated with the NEBNext microbiome DNA enrichment kit (New England Biolabs, Ipswich, MA, USA) for human DNA removal, before an additional purification using AMPure XP SPRI beads and final elution in 15μl TE buffer. Samples were extracted in batches, with a negative control of sterile 0.9% saline prepared alongside each batch using this same protocol.

### Library preparation and Illumina MiSeq sequencing

DNA extracts quantified as ≥0.2ng/μl were sequenced on a MiSeq desktop sequencer (Illumina, San Diego, CA, USA). Libraries were prepared as previously described, using a variation of the Illumina Nextera XT protocol (34). Briefly, 1ng of DNA was prepared for sequencing following the Illumina Nextera XT protocol, with the modification of 15 cycles during the index PCR. Libraries were quantified using a Qubit 2.0 fluorometer and their average size determined with an Agilent 2200 Tapestation (Agilent Technologies, Santa Clara, CA, USA) before being manually normalised. Libraries were prepared and sequenced together in the same batch. Paired-end sequencing was performed using a 600-cycle MiSeq reagent kit v3 and samples were sequenced in batches of between 1 and 13 on a single flow cell.

### Bioinformatics analysis

Raw sequencing reads were adapter trimmed using BBDuk (https://sourceforge.net/projects/bbmap/) with the following parameters: minlength=36, k=19, ktrim=r, hdist=1, mink=12 and the adapter sequence file provided within the bbmap package. Taxonomic classification of trimmed reads was performed using Kraken (35) and a bespoke database constructed from all bacterial genomes deposited in the NCBI RefSeq database as of January 2015 (updated January 2017 for validation set, see below), with default parameters and no K-mer removals. Where no refseq genome was available for an organism cultured from a PJI at OUH since June 2013, available whole-genome assemblies were also added to the database where available in NCBI. Additionally, the human genome GRCh38 was included in the database to allow detection of host DNA. An optimum filtration threshold, using Kraken-filter, that balanced false-positive removal and sensitivity was determined using simulated datasets of reference genomes. Reference genomes representative of common pathogenic species were used to generate simulated Illumina MiSeq datasets and analysed with Kraken using different filtration thresholds. A threshold value of 0.15 provided optimum read classification sensitivity whilst minimizing spurious results. Kraken output was visualised using Krona (36).

### Statistical analysis

In order to correct for samples which may contain small numbers of contaminating and non-specific bacterial reads, a threshold was determined to identify the presence of true infection, using the first 50 samples sequenced as a derivation set. Two thresholds (1 and 2), and three parameters (a-c), were used to determine true infection: 1) samples with more reads from a given species than an upper read cut-off (a) were included; 2) samples with more species-specific reads than a lower read cut-off (b) and with the percentage of species-specific reads as a proportion of all bacterial reads present above a percentage cut-off (c) were also included. Parameter values were selected by numerical optimisation, using R version 3.3.2, comparing sequencing results to sonication fluid culture results, and maximising the value of the Youden Index (37) (sensitivity + specificity - 1). Sensitivity was calculated taking each species identified from each culture-positive sonication sample as a separate data point. Specificity was calculated using the total number of sonication samples as the denominator; as such samples contaminated by more than one species were counted as one false-positive.

To ensure that read cut-off parameters were chosen without a penalty for potentially difficult to culture anaerobic species, the specificity value optimised was adjusted. Potential ‘false-positive’ sequencing results with plausible fastidious anaerobic causes of infection (including *Fusobacterium nucleatum, Propionibacterium acnes* and *Veillonella parvula)* in culture-negative samples were excluded when calculating the specificity value used for parameter optimisation.

Where bacterial reads were detected over the thresholds described above in a negative control, that sample was deemed to be contaminated. In the derivation set, in order to maximise the number of sequences available for analysis, only samples with evidence of the same contaminating organisms were excluded from each contaminated batch, rather than discarding the whole batch. During the derivation phase of the study, several batches of samples were found to be contaminated with DNA from other studies performed concurrently in the same research laboratory. Six of eight saline negative control extracts displayed contamination with a single or multiple species at read numbers exceeding the determined diagnostic thresholds. All samples within these batches that displayed similar contamination were excluded from subsequent analysis if Kraken classification resulted in >100 reads corresponding to the majority of the contaminating species. A total of 22 samples (in addition to the 50 successfully sequenced) were excluded on this basis (Figure 1). In batches 4 and 5 the negative controls were contaminated with *Staphylococcus aureus, Escherichia coli*, and *P. acnes*, and 15 samples were excluded with >100 reads from ≥ 2/3 species; in batch 6 the negative control was contaminated with *Serratia marcescens, Klebsiella pneumoniae, E. coli*, and *P. acnes*, and 2 samples with >100 reads from ≥3/4 species were excluded; in batches 2, 9 and 10 the negative control was contaminated with *P. acnes*, and 5 samples were excluded with >100 *P. acnes* reads. To address this issue, prior to the validation phase of the study, all pipettes, laminar flow and PCR hoods and laboratory benches used for DNA extraction and library preparation were deep-cleaned with Virkon disinfectant and RNase AWAY surface decontaminant (Thermo Fisher Scientific, Waltham, MA, USA) in order to remove any possible sources of microbial or DNA contamination. All DNA extraction and library preparation reagents were replaced, and used in pre-prepared per-batch aliquots used exclusively for this study. Sonication fluid samples were handled one at a time in the laminar flow hood, which was cleaned as above between each sample. Fresh gloves were worn each time a new sample was handled during the DNA extraction phase of the protocol. Having implemented these changes, for the validation phase, a more stringent quality control standard was applied, requiring the negative control to be contamination-free for any of the samples in a batch to be analyzed.

**Figure 1.**
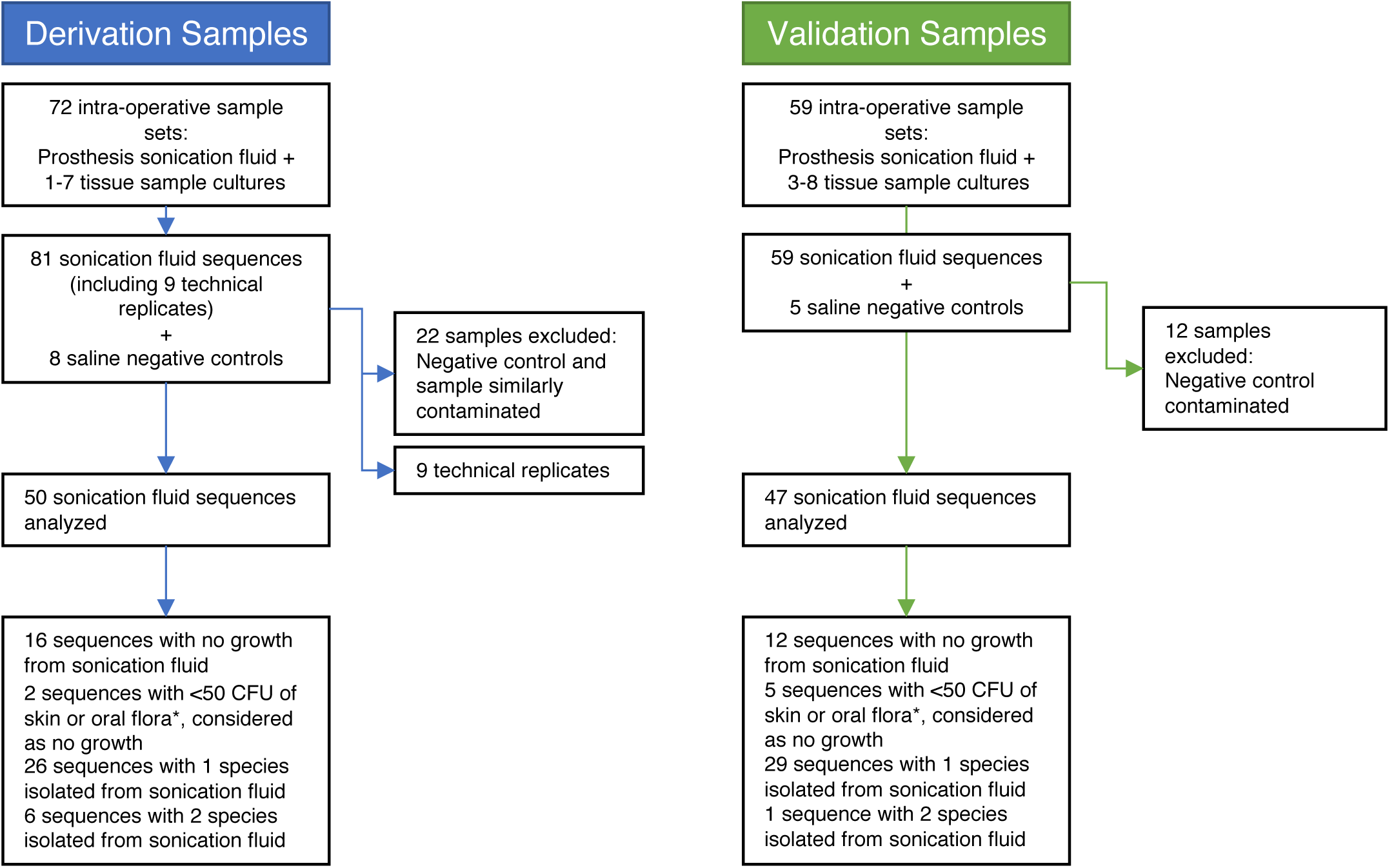
Study samples and quality control. *\*StaphyIococcus epidermidis*, other coagulase-negative Staphylococci, viridans group Streptococci, *Propionibacterium acnes*

### Technical replicates

To ensure sequencing reproducibility one DNA sample was sequenced twice and biological replicates (DNA extraction process repeated) were sequenced for six samples (four in duplicate and two in triplicate). Samples extracted and sequenced as replicates showed good reproducibility. In four duplicate and one triplicate culture-positive samples the same species was recovered by WGS on all occasions (samples 164, 171, 182, 183, and 193). A single replicate, 182a, had an additional, likely contaminating, species identified (not found in sonication fluid or PPT culture). A single culture-negative sample (176) was processed in triplicate. One of the three replicates (176a) had an apparent contaminating species identified (also not found in sonication fluid or PPT culture).

## Results

A total of 131 sonication fluids from patients undergoing revision arthroplasty or removal of other orthopaedic devices, were aerobically and anaerobically cultured and underwent metagenomic sequencing (Figure 1). Additionally, a median (IQR) [range] 5 (4–5) [1-8] PPT samples were cultured from each patient. *S. aureus*, isolated from 22% of sonication fluids and 29% PPTs, and *Staphylococcus epidermidis*, from 16% sonication fluids and 25% of PPTs, were the 2 most frequently cultured species (Table 1).

**Table 1:** Summary of species observed in microbiology culture for sonication fluid and periprosthetic tissue (PPT), presented by joint/implant type. Results reported as number observed in sonication fluid (number observed in PPT). ^a^Metalwork comprises plates and/or screws from tibia (*n*=3), femur (*n*=4), spine (*n*=2), foot (*n*=2), humerus (*n*=1), ankle (*n*=1) and ulna (*n*=1).

| Species | Ankle<br>$n=6$ | Hip<br>$n=32$ | Knee<br>$n=42$ | Metalwork<sup>a</sup><br>$n=14$ | Shoulder<br>$n=3$ | Total<br>$n=97$ |
| --- | --- | --- | --- | --- | --- | --- |
| Staphylococci |  |  |  |  |  |  |
| *Staphylococcus aureus* | 0 (1) | 5 (9) | 10 (11) | 5 (6) | 1 (1) | 21 (28) |
| *Staphylococcus condimenti* |  |  | 1 (0) |  |  | 1 (0) |
| *Staphylococcus epidermidis* | 1 (1) | 6 (10) | 9 (12) | 0 (1) | 0 (1) | 16 (25) |
| *Staphylococcus lugdunensis* | 0 (1) | 1 (1) |  | 1 (1) |  | 2 (3) |
| Coagulase-negative *Staphylococcus* | 1 (0) | 0 (3) | 1 (2) | 0 (1) | 1 (0) | 3 (6) |
| Streptococci |  |  |  |  |  |  |
| *Streptococcus agalactiae* |  |  | 2 (2) |  |  | 2 (2) |
| *Streptococcus dysgalactiae* |  | 0 (1) |  |  |  | 0 (1) |
| *Streptococcus oralis* |  | 0 (2) |  |  |  | 0 (2) |
| *Streptococcus pneumoniae* |  | 1 (1) |  |  |  | 1 (1) |
| *Streptococcus vestibularis* |  |  |  | 1 (0) |  | 1 (0) |
| Enterococci |  |  |  |  |  |  |
| *Enterococcus faecalis* |  | 3 (4) | 2 (2) | 0 (1) |  | 5 (7) |
| *Enterococcus faecium* |  | 0 (1) | 3 (3) |  |  | 3 (4) |
| Enterobacteriaceae |  |  |  |  |  |  |
| *Citrobacter koseri* |  | 1 (1) |  |  |  | 1 (1) |
| *Citrobacter species* |  |  | 1 (1) |  |  | 1 (1) |
| *Enterobacter cloacae* | 1 (1) |  | 1 (1) |  |  | 2 (2) |
| *Escherichia coli* |  | 1 (2) |  |  |  | 1 (2) |
| *Klebsiella oxytoca* |  |  |  | 0 (1) |  | 0 (1) |
| *Klebsiella pneumoniae* |  |  | 1 (1) |  |  | 1 (1) |
| *Morganella morganii* |  |  | 2 (2) |  |  | 2 (2) |
| *Proteus mirabilis* |  | 0 (1) | 1 (1) |  |  | 1 (2) |
| *Serratia marcescens* |  |  | 1 (1) |  |  | 1 (1) |
| Corynebacteria |  |  |  |  |  |  |
| *Corynebacterium amycolatum* |  | 0 (1) |  |  |  | 0 (1) |
| *Corynebacterium aurimucosum* |  | 0 (1) |  |  |  | 0 (1) |
| *Corynebacterium propinquum* |  |  |  | 0 (1) |  | 0 (1) |
| *Corynebacterium striatum* |  | 0 (3) |  | 0 (1) |  | 0 (4) |
| Other |  |  |  |  |  |  |
| *Aeromonas* species |  |  |  | 0 (1) |  | 0 (1) |
| *Aeromonas hydrophila* |  |  |  | 1 (0) |  | 1 (0) |
| *Arcanobacterium haemolyticum* |  | 1 (0) |  |  |  | 1 (0) |
| *Bacillus* species | 0 (1) | 1 (2) |  |  |  | 1 (3) |
| *Fingoldia magna* |  |  |  | 1 (0) |  | 1 (0) |
| *Gemella morbillorum* |  | 1 (1) | 1 (1) |  |  | 2 (2) |
| *Granulicatella adiacens* |  |  |  | 0 (1) |  | 0 (1) |
| *Micrococcus luteus* |  |  |  | 0 (1) |  | 0 (1) |
| *Mycobacterium fortuitum* |  |  | 0 (1) |  |  | 0 (1) |
| *Propionibacterium acnes* |  | 1 (1) | 2 (0) |  |  | 3 (1) |
| *Propionibacterium* spp. |  | 0 (1) |  |  |  | 0 (1) |
| *Pseudomonas aeruginosa* | 1 (2) | 0 (2) |  | 1 (2) |  | 2 (6) |
| No growth | 3 (3) | 12 (7) | 7 (6) | 5 (4) | 1 (1) | 28 (21) |
| **Total** | **7 (10)** | **34 (55)** | **45 (47)** | **15 (22)** | **3 (3)** | **104 (137)** |

From the first 72 sonication fluid samples sequenced 22 samples from six batches were excluded, as these samples and negative controls from the same batches showed similar contamination (see Methods, Figure 1). The remaining 50 samples, the derivation set, were used to determine optimal sequence thresholds for identifying true infection. Of 59 subsequently sequenced validation samples, 12 from a single batch were excluded as the negative control was contaminated with *P. acnes*, leaving 47 validation samples sequenced in batches with uncontaminated negative controls.

The 97 sonication fluid samples passing sequencing quality control checks were obtained predominantly from knee (42/97, 43%) and hip (32, 33%) PJI, with other samples from ankle (6, 6%), and shoulder (3, 3%) PJI, and other orthopaedic device infection (14, 14%) (Supplementary Table 2). The median (IQR) [range] sonication fluid volume was 200ml (100-400ml) [15-400ml] (Supplementary Table 2). On culture, 35 (36%) sonication fluid samples had no growth, or less than <50 CFU of an organism not considered to be highly pathogenic (skin and oral flora), 55 (57%) had a single organism isolated, and 7 (7%) two organisms isolated. Greater than 10^6^ reads were achieved in 91/97 (94%) samples. Taxonomic classification by Kraken identified a median (IQR) [range] 0.72% (0.01-0.41%) [<0.01% - 24.0%] of reads as bacterial, with <1% bacterial reads in 84/97 (87%) samples. Human reads accounted for >90% of reads in 94/97 (97%) of samples. Six test samples were processed with and without the NEBNext microbiome DNA enrichment kit. Use of the kit did not reduce the amount of human DNA sequenced. The mean proportion of reads classified as human with the enrichment kit was 98.4%, and without it 98.2% (p=0.06, Supplementary Table 1).

Optimal thresholds for determining if samples contain low-level contamination or true infection were determined by numerical optimisation, choosing thresholds that maximised the sensitivity and specificity of sequencing (Figure 2). The final thresholds chosen to determine the presence of true infection were ≥1150 reads from a single species, or ≥125 reads from a single species if ≥15% of the total bacterial reads also belong to that same species.

**Figure 2.**
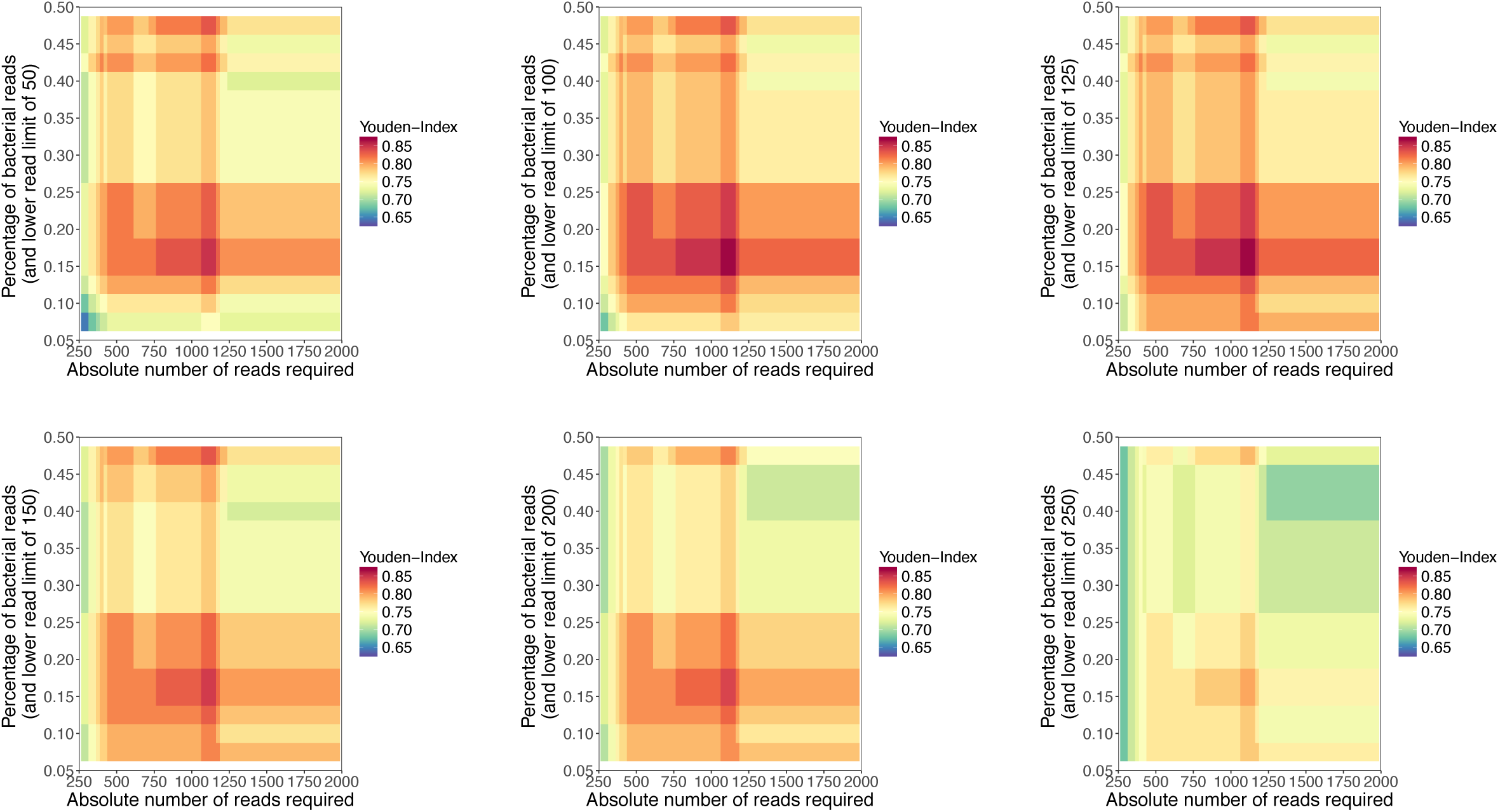
Sequencing data filtering calibration heatmaps. Two thresholds (1 and 2), and three parameters (a-c), were used to determine true infection: 1) samples with more reads from a given species than an upper read cut-off (a, x-axis) were included; 2) samples with more species-specific reads than a lower read cut-off (b, panels) and with the percentage of species-specific reads as a proportion of all bacterial reads present above a percentage cut-off (c, y-axis) were also included. Youden-Index = (sensitivity + specificity) - 1.

Table 2 compares sonication culture results with metagenomic sequencing findings applying these thresholds. PPT culture results, and the consensus microbiology diagnosis based on both sonication and PPT samples are also given for comparison. Compared to sonication fluid culture, metagenomic sequencing had an overall species-level sensitivity of 61/69 (88%, 95%CI, 77-94%). Sensitivity in the derivation samples was 35/38 (92%, 79- 98%), and in the validation samples 26/31 (84%, 66-95%). Three samples were identified to the genus level only. Hence overall genus-level sensitivity was 64/69 (93%, 84-98%). Of the other five samples where the species cultured was not identified on sequencing, two samples cultured a coagulase-negative *Staphylococcus*, not identified on tissue culture, one sample was polymicrobial (where several species found in sonication fluid or tissue were identified, but not all) and the remaining two samples failed to detect a pathogen found in sonication and tissue fluid.

**Table 2:**
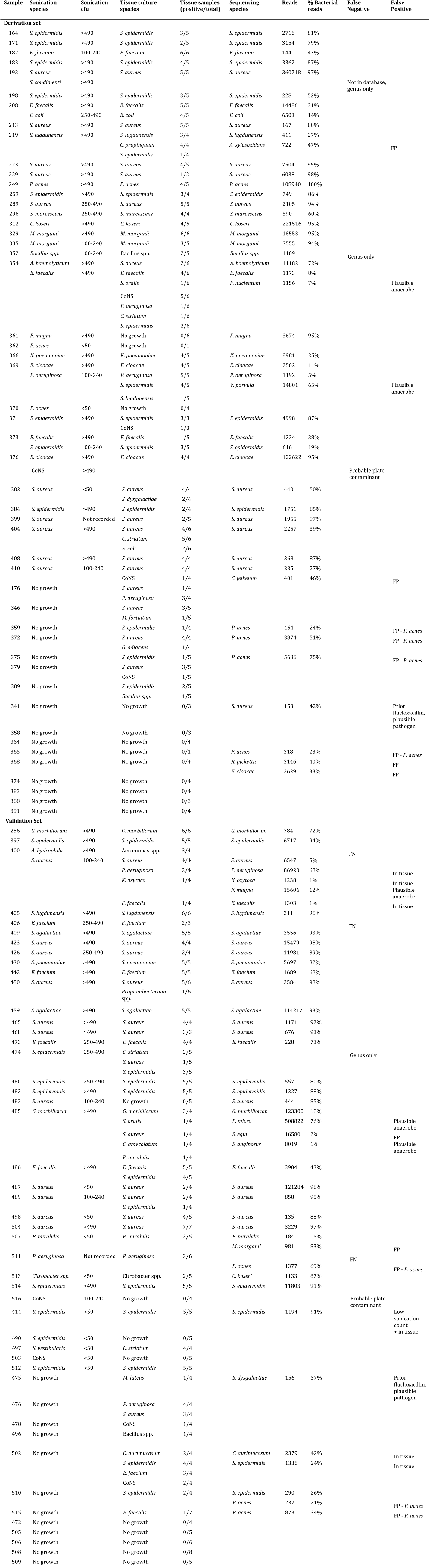
Comparison of species identified from sonication fluid and PPT culture with species identified from metagenomics sequencing reads for all samples passing thresholds for analysis in the derivation (*n*=50) and validation (*n*=49) data sets. CoNS, coagulase-negative *Staphylococcus* species; FN, false-negative result; FP, false-positive result. See Supplementary Table 2 for genus details.

Overall species level specificity was 78/97 (80%,71-88%). However, of 19 samples where additional species were identified on sequencing compared to sonication culture, three had the same species found in tissue culture (but not in sonication fluid, or at <50 CFU; samples 400, 414, 502). Four samples had plausible anaerobic causes of infection (*Fusobacterium nucleatum, Veillonella parvula, Finegoldia magna, Parvimonas micra* [identified alongside *Streptococcus anginosus*]) (samples 354, 369, 400, 485). Samples 341 and 475 contained *S. aureus* and *Streptococcus dysgalactiae* DNA respectively, both in patients who had received prior flucloxacillin and no microbiological diagnosis was reached based on culture. However, 12 samples (including sample 485) had other species found on sequencing not otherwise identified. In some cases these were clearly laboratory contaminants, e.g. sample 219 contained *Achromobacter xylosoxidans* reads and an *A. xylosoxidans* culture-positive sample was sequenced in the same batch from a concurrent study. Notably *P. acnes* was a common contaminant occurring in 7/97 (7%) samples overall. Adjusting for plausible fastidious causes of infection, species found in concurrently obtained PPT samples, and prior antibiotics, i.e. assuming these samples were actually genuinely positive for the species found on sequencing, overall species-level specificity was 85/97 (88%,79-93%), 43/50 (86%,73-94%) in the derivation samples and 42/47 (89%,77-96%) in the validation samples.

Figure 3 shows the relationship between the proportion of sequence reads obtained that were classified as bacterial, the sonication fluid culture CFU counts, and the concordance between sonication fluid culture and sequencing. Sequencing false positive results were more likely when cultures were negative.

**Figure 3.**
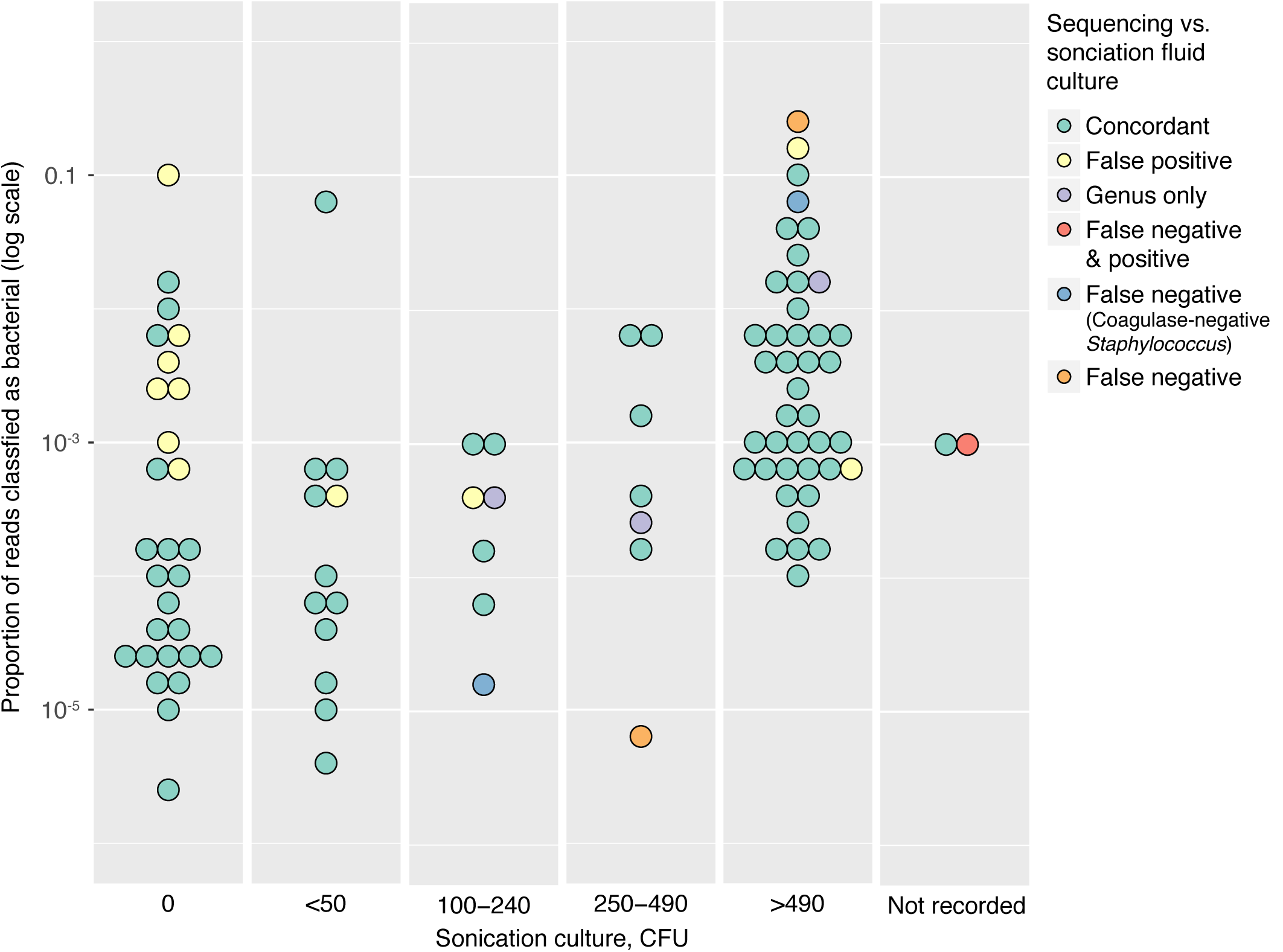
Sonication culture and sequencing comparison. The proportion of sequencing reads classified as bacterial is shown on the y-axis on a log scale, and the number of CFU from sonication fluid culture on the x-axis. Markers are coloured by the concordance of sonication fluid culture and sequencing. A single marker is shown per patient sample, where only one of several species isolated was found by sequencing this is shown as a false negative marker, similarly any sample with one or more false positive species identified by sequencing is shown as false positive. False negative results where a coagulase-negative *Staphylococcus* was cultured from sonication fluid, but not found in tissue samples or on sequencing are shown separately, as are samples only identified to the genus level by sequencing. Results were very similar if absolute numbers of bacterial reads were plotted on the y-axis instead.

More simplistic thresholds for determining true infection performed less well. Within the derivation set, using a single cut-off for the proportion of bacterial reads from a given species, irrespective of the absolute numbers of bacterial reads present, the optimal cut-off value was 25%. Using this threshold, species-level sensitivity was 30/38 (79%) and adjusted specificity 44/50 (88%). Similarly, if only a single absolute read number cut-off is used the optimal value is 410 reads from a single species, sensitivity 30/38 (79%) and adjusted specificity 45/50 (90%).

Sequencing results were also compared to a consensus microbiology diagnosis based on IDSA guidelines (4), considering any species isolated twice or any virulent species isolated as a cause of infection, combining sonication and PPT culture results (Supplementary Table 2). 65/97 (67%) samples showed complete agreement between the consensus species list from culture and sequencing, 15/97 (15%) a partial match with at least one species found on culture also found on sequencing, 15/97 (15%) had none of the species cultured found on sequencing, and 2/97 (2%) had a plausible additional species found on sequencing not found on culture.

## Discussion

Diagnosis of PJI by culture of sonication fluid and PPT is not always conclusive, and may take up to 10-14 days for slow-growing organisms. Here we assess, for the first time, the use of metagenomic sequencing of complete bacterial DNA extracts obtained direct from sonication fluid in the diagnosis of PJI. We develop a novel filtering strategy to ensure that low-level contaminating DNA is successfully ignored, while infections are detected accurately. Compared to sonication fluid culture, metagenomic sequencing achieved a species-level sensitivity of 88% and specificity of 87% after adjusting for plausible fastidious causes of infection, species found in concurrently obtained PPT samples, and prior antibiotics. Importantly we demonstrate similar performance of our method and filtering algorithm in the subset of samples that formed an independent validation set, sensitivity 84%, adjusted specificity 89%.

Sequencing failed to identify an organism cultured from sonication fluid for eight samples. For two samples a coagulase-negative *Staphylococcus* was cultured, but only from sonication fluid and not from tissue samples. These isolates therefore could plausibly have been plate contaminants, and not present in the DNA sequenced. For three other samples identification to the genus-level was possible. One sample contained *Staphylococcus condimenti* which was not included in our custom Kraken database, highlighting that despite including 2786 bacterial genomes this approach is only as good as the database that is used. Another sample was identified as a *Bacillus spp.* both on culture and by sequencing, and the third by sequencing as *Staphylococcus spp.* in the context of a mixed *Staphylococcus* infection. For the three remaining samples, sequencing failed to identify a pathogen found on culture.

Sequencing was also able to detect potential pathogens not identified by sonication fluid culture. For three samples we identified additional species from sequencing that were supported by the tissue culture findings, suggesting in some settings sequencing may be more sensitive that sonication fluid culture alone without PPT culture. We also identified four examples of probable anaerobic pathogens not identified by routine anaerobic culture of sonication fluid or PPT: *Fusobacterium nucleatum, Veillonella parvula, Finegoldia magna, Parvimonas micra*, and we were also able to identify a plausible pathogen in two patients who had received prior antibiotics where the routine microbiology was uninformative.

Controlling for contamination during sampling and culture is a major challenge in investigating PJI, and underlies why multiple independent PPT samples remains the gold standard for diagnosis. Contamination is an even greater concern in molecular diagnostics given the additional potential for DNA contamination. There are published examples demonstrating the potential for contamination leading to misinterpretation of sequencing data from clinical specimens (38, 39). In our laboratory samples were handled in laminar flow hoods and extracted in a dedicated pre-PCR extraction laboratory. DNA was handled in a PCR hood and sequencing libraries were manipulated in a dedicated post-PCR sequencing laboratory. Despite these measures, we still observed contamination in some of our samples. During the derivation phase of our study it is likely that one or more of the reagents used became contaminated with DNA from other sequencing projects in our laboratory. Although we were able to account for this in our analysis, and then validate our findings in a separate set of samples having resolved this issue, it demonstrates that rigorous laboratory practice would be key to deploying our method. There may also be a role for sealed systems that perform DNA extraction and sequencing in a separated environment. Our experience also re-enforces the requirement that negative controls are included in each sequencing batch, as is routine in molecular microbiology diagnostic assays, to ensure contamination is detected if it does occur. A limitation of our study is that the saline used for sonication was not PCR-grade, and this could be considered in future work.

Despite addressing the major contamination issues in the derivation phase, contamination with *P. acnes* remained an issue in one of our validation batches. Overall, false positive results for *P. acnes* were found in 7% of samples. Species-specific filtering may be required to address this, our one true-positive sample with *P. acnes* present on culture had >10^5^ *P. acnes* reads. However, larger datasets are required than ours to address this definitively. In the mean-time, even with molecular diagnostics, the value of multiple samples per patient remains.

Sonication fluid can be a large volume sample, typically 50-400ml. As a result, the microbial cells released from the orthopaedic device during sonication are likely to be heavily diluted. This, coupled with the simultaneous release of any human cells from the prosthesis and transfer of blood along with the device, results in a sonication fluid sample that is both low in bacterial cells and high in contaminating host cells. An effective microbial DNA extraction protocol is necessary to isolate as much bacterial DNA as possible, while limiting the amount of host DNA in the final extract. Our results demonstrate that despite efforts to filter out human cells, or remove human DNA post-extraction, host DNA accounted for >90% of reads in the majority of samples sequenced. Use of a specialist microbiome enrichment kit did not improve bacterial DNA yield. However, if the efficiency of human DNA removal can be improved in future this might significantly add to the precision of metagenomic sequencing as more sequencing effort would be appropriately directed towards potential pathogens.

In addition to the issues around contamination with bacterial and human DNA, a further limitation of our study as designed is that it undertakes a laboratory-level comparison of sonication fluid culture and metagenomics sequencing. We did not have ethical approval to review patient notes and so were unable to compare sonication fluid sequencing to the presence of a final overall diagnosis of infection. Future studies should consider how sequencing might contribute to the overall diagnosis of PJI, as part of assessment that jointly considers clinical, histological, and microbiological data.

This study demonstrates as a proof of principle that metagenomic sequencing can be used in the culture-free diagnosis of PJI directly from sonication fluid. Improvements to the method of human DNA removal from direct samples before sequencing are ongoing, and if these are successful, this is likely to greatly improve the efficiency, and therefore accuracy, of metagenomic sequencing. Generating greater numbers of bacterial reads direct from clinical specimens may make prediction of antimicrobial susceptibilities direct from samples possible, as has been achieved from whole-genome sequencing of cultured organisms (25–28). If this can be achieved reliably, it is possible that sequencing can offer a complete microbiology diagnosis without the need for culture. The increasing availability of portable, rapid, random-access strand sequencing technology offers the potential that in future sequencing may become a same-day diagnostic tool. Applications of rapid sequencing in PJI might include perioperative microbiological diagnosis to guide local intraoperative antimicrobials, for example in cement or beads. Earlier diagnosis may also ensure post-operative antimicrobials are more focused, improving antimicrobial stewardship, while treating resistant organisms effectively. Earlier diagnosis may also reduce hospital stays and therefore reduce costs. Sequencing is also likely to be helpful in situations where multiple samples containing the same commensal species are identified. Sequencing will be able to determine whether these are clonal, suggesting true infection rather than contamination, instead of having to rely on current proxies such as antibiograms, which only imperfectly distinguish non-clonal isolates. Ultimately, same-day sequencing may significantly improve the precision, efficiency and cost of PJI care. This study provides a foundation for further development towards this goal.

## Acknowledgements

The authors thank the microbiology laboratory staff of the John Radcliffe Hospital, Oxford University Hospitals NHS Foundation Trust, for providing assistance with sample collection and processing.

